# Deciphering Hematopoiesis at single cell level through the lens of reduced dimensions

**DOI:** 10.1101/2022.06.07.495099

**Authors:** Prashant Singh, Yanan Zhai

## Abstract

Hematopoiesis plays a critical role in maintaining a diverse pool of blood cells throughout human life. Despite recent efforts with single-cell data analyses, the nature of the early cell fate decisions and compartmentalization of progenitors remains contentious due to the sparsity and noise of the data. Using publically available single-cell RNA-Seq hematopoietic data from bone marrow, with three different matrix factorization approaches to recover associated gene modules from cell clusters reveals a tri-directional and hierarchically-structured transcriptional landscape of hematopoietic differentiation. We also devised a bootstrap method, which in combination with the above can better characterize the progenitor compartments and retrieve cellular hierarchies.

## Introduction

Hematopoiesis is the formation of cellular components of blood and it consists of an organized set of developmental stages from a hematopoietic stem or progenitor cells (HSPCs) that then differentiate into more mature hematopoietic lineages (e.g-Monocytes, B and T-cells, etc.), which then provide all the key functions of the immune system^1–4^. Complex biological processes (e.g-Transcriptional dynamics) behind hematopoiesis can help clinicians to better understand the processes responsible for blood disorders and cancers like leukemia. Furthermore, hematopoietic stem or progenitor cells (HSPCs) can be used as a model system for understanding tissue stem cells and their role in aging and oncogenesis^3,5^. To access the cellular hierarchy of hematopoiesis and its transcriptional heterogeneity, comprehensive profiling and clustering using single-cell RNA sequencing (scRNAseq) have been performed previously^6–9^. These studies readily detected cellular subsets with lineage-specific gene expression, including putative erythroid, monocytic, and granulocytic lineage-restricted progenitors. However, distinguishing between the previously reported multipotent and oligopotent cell types, such as human multipotent progenitors (MPPs), common lymphoid and myeloid progenitors (CLPs and CMPs) is still complicated. The reason for this complexity is probably ultra-high dimensionality and sparsity of the single-cell datasets. scRNA-seq datasets are highly sparse and contain signals from gene expressions at the single-cell level that control biological systems and may characterize complex biological processes or programs (CBPs). To reveal low-dimensional structures from high-dimensional data, matrix factorization (MF) techniques are regularly used to reflect on those CBPs. These techniques can unravel new biological insights from various scRNA-seq datasets in applications ranging from pathway discovery, cellular hierarchies to timecourse analysis^10^. In this work, we applied three popular MF techniques (Principle component analysis; PCA, Independent component analysis; ICA, and Non-negative matrix factorization; NMF) as a primary dimension reduction on a publicly available scRNA-seq dataset to establish cellular hierarchies in hematopoiesis and better characterize progenitor compartments based on differential gene modules or programs obtained. We also applied bootstrapping of cell clusters in each case to assess the cluster stability and their interconnections or hierarchies. We believe that this work could serve as a reference map for healthy hematopoiesis and a guideline to pick an effective MF strategy in combination with a bootstrapping for new scRNA-seq datasets.

## Results

### PCA captures dominant sources of varitation but fails to distinguish among progenitor compartments

We performed PCA on bone-marrow-derived single-cell transcriptomic data (four healthy controls, ~21000 cells, from GSE139369), followed by uniform manifold approximations (UMAP)^11^, and this allowed us to recover variations in expression signals from most of the clusters (Figure 1A, B). Low dimensional structure of the data (Figure 1B) suggest the tri-directional developmental trajectory i.e. from is HSPCs to Erythroids via Megakaryocyte–erythroid progenitor cells (MEPs), HSPCs to differentiated myeloid lineages (such as CD14^+^monocytes, Dendritic Cells, etc.), and HSPCs to differentiated lymphoid lineages like B and T-cells (Fig1B). For mature blood cell types, we identified clear transcriptional signatures of erythroid cells (expressing HBB, HBA1, etc.), monocytes (MoCs expressing CD14, FCN1, VCAN, etc.), plasmacytoid dendritic cells, and Dendritic cells (pDCs and DCs; expressing LILRA4, IRF7/8, HLAs, etc.) with an additional cluster of highly cycling DCs (cDCs). Committed progenitors like granulocyte-monocyte progenitors (GMPs) cells (expressing AZU1, MPO, and PRTN3) were also clearly distinguishable (Fig1A, B). Although granulocyte progenitors were present in our dataset, we could not trace neutrophils, basophils, and eosinophils. In the lymphoid compartment, we identified T-cells and NKT cells (expressing CD3E, GNLY, NKG7, etc.) and mature B cells (expressing MS4A1 and IGHD) (Figure 1A, B). The B cell lineage also included plasma-B cells (pBCs), which showed expression of CD79A/B, and VPREB1/3 (Figure 1A, B).

**Fig. 1.**
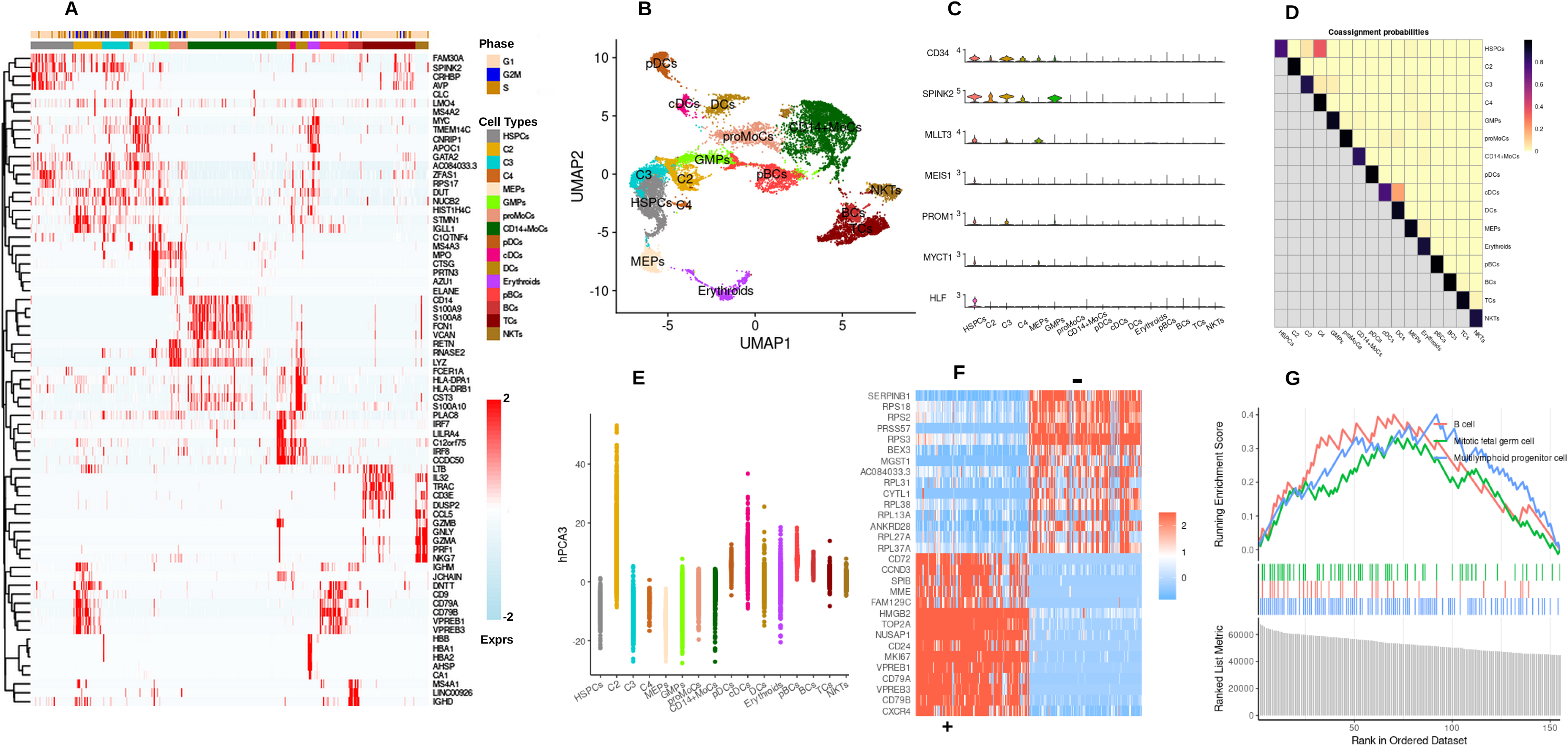
Exploration of Bone marrow SC-data with PCA. **A,** Heatmap representing the top genes, differentially expressed among various cell types. Colored annotation bars represent the cell cycle phase of each cell and cell types classifications. Colored Scales represent a normalized expression. **B, UMAP** representation of each cell class. **C, Violinplot** showing expression of prominent HSC markers among all cell types. **D, Heatmap** of coassignmnet probabilities of multiple bootstrap sampling and clustering. **E, Plot** revealing harmony corrected principal component 3 (hPCA3) explains variation for cluster 2 (C2). **F, G, Top gene modules** positively and negatively associated with C2 and GSEA on those modules indicating probable cell classification of C2.

A cluster of HSPCs showed high expression of CD34, SPINK2 genes which play a crucial role in role in early hematopoiesis maintenance^12,13^. But these markers are equally expressed in other nearby clusters (C2, C3, and C4)(Fig1A,C). Hence, we used markers like MLLT3, a crucial regulator of human HSC maintenance, HLF, a transcription factor (TF) involved in preserving quiescence in HSCs, MEIS1, a TF involved in limiting oxidative stress in HSCs, which is necessary for quiescence, and MYCT1, a gene involved in the regulation of hematopoietic differentiation^14-20^, that showed differential expression among these progenitor compartments (Fig1C) and helped us for annotating HSPCs. Also, majority of cells in that cluster belong to the G1 phase of the cell cycle (Fig1A), which also suggests the presence of quiescent HSCs^21^. Other compartments (C2, C3, and C4) could be CLPs, CMPs, or MPPs but annotating them with confidence with gene programs retrieved by PCA is challenging. A bootstrap sampling and clustering showed overlaps of assignment probabilities between HSPCs with C3 and C4 but not with C2 (Fig1D) indicating C2 might be a different class of progenitors. Next, we tried to identify which principal component is explaining the variations for C2 and identified the third component (PCA3) is responsible (Fig1E). Top gene modules positively associated with this component (Fig1F) indicates lymphoid-specific commitments (e.g-SPIB) and early B-cell development signals (e.g-VPREB1), suggest C2 might be CLPs (expressing signals like MME, CD24, CD72). Gene set enrichment analysis (GSEA)^22^ with ranked gene programs obtained by PCA3 with CellMarker database^23^ also depicted enrichment terms related to B-cell and early lymphoid development (Fig1G).

Further, to determine progenitor classes of C3 and C4, we also tried to find principal components (PCs) linked with C3 and C4 but failed. We also attempted to perform gene get enrichment analysis (GSEA) against using DE gene modules from C2, C3, and C4 but no terms were found enriched under the *p-value* cutoff of 0.05, suggesting that gene modules retrieved from PCA were not able to distinguish among these progenitors.

### ICA infers additive patterns in data

In contrast to PCA, the ICA components cannot be ranked by the percent variation explained. Instead, ICA tends to find statistically independent sources of variation in the data^24^. After performing ICA followed by UMAP on the same SC-data, we again retrieved tri-directional trajectory from HSPCs (Fig2A, B). Similar to PCA results, we observed easier annotations of differentiated cell types, e.g-Erythroids expressing HBB, HBA1, etc., and B-Cells expressing MS4A1. Although more cell types were recovered, e.g-Naive T-cells (Tn; expressing IL7R) maturing towards T-cells and NKTs, and CD16^+^Monocytes (expressing a strong marker like FCGR3A) (Fig2A, B). Unlike PCA results, we observed a slightly better separation among progenitor compartments as CLPs are distinguishable with higher expression of DNTT and RAG1 (markers for lymphoid commitment)^25^. Nevertheless, clusters 2 and 3 (C2, C3) were still quite intermixed with other progenitor clusters and were difficult to classify (Fig2A, B). The vicinity of C3 with other myeloid lineages (like GMPs) (Fig2B) indicated that it could be CMPs, and bootstrap overlaps of coassignment probabilities among C3, GMPs, and promonocytes also hinted the same (Fig2C). GSEA with ranked differentially expressed (DE) gene modules (~160 genes) obtained from C3 (Fig2D) showed monocyte progenitor related terms, further confirming the possibility of C3 being CMPs. C2 might be MPP as bootstrapping shows the overlapping co-assignment probabilities with CLPs, CMPs, and MEPs, but we couldn’t retrieve any component or gene modules exclusively associated with C2, which could be used for GSEA. Interestingly we also found ICA29’s association with HSPCs and few of the linked gene programs (Fig2E, F) retrieved (e.g-MLLT3, HLF, MYCT1, and HOPX) were already known for their role in hematopoietic differentiation and maintenance^14,15,20,26^. Altogether, these results indicate ICA infers gene programs that are additive in nature and better to define cellular hierarchies among multiple cell types, which could be simplified and viewed with bootstrapping the clusters.

**Fig. 2.**
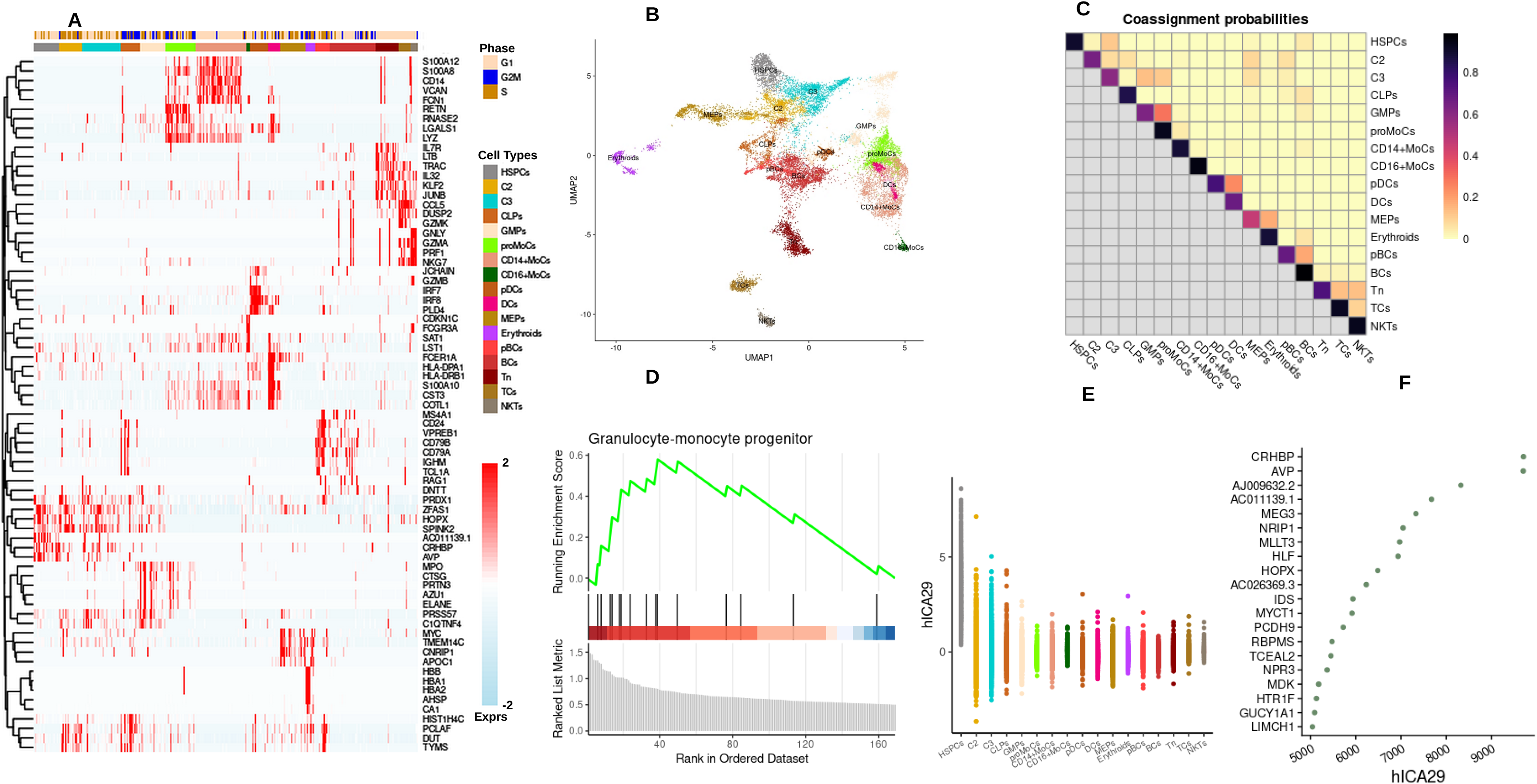
Exploration of Bone marrow SC-data with ICA. **A,** Heatmap representing the top genes, differentially expressed among various cell types. Colored annotation bars represent the cell cycle phase of each cell and cell types classifications. Colored Scales represent a normalized expression. **B, UMAP** representation of each cell class. **C, Heatmap** of coassignmnet probabilities of multiple bootstrap sampling and clustering. **D, GSEA** on C3 using DE gene modules. **E, Plot** revealing harmony corrected independent component 29 (hICA29) separates HSPCs from other. **F, Top gene** programmes (ranked) linked with HSPCs.

### NMF learns distinct patterns from the data

Similar to ICA, NMF can also not be ranked by the percent variation explained but it has the ability to automatically extract sparse and easily interpretable non-negative factors in an unsupervised manner^10,27^. NMF followed by UMAP on SC-data also showed a tri-directional developmental trajectory of blood cells, and smooth classification of differentiated cell types (Fig3A, B), similar to ICA or PCA. Interestingly bootstrap clustering displayed strictly nonoverlapping coassignment probabilities (Fig3C), indicating NMF based clustering might be retrieving idiosyncratic gene modules and could be useful to classify progenitor compartments. As we were unable to classify MPPs (with confidence) in previous cases, we tried to find factors associated with MPPs. We found gene modules representing the twenty-fifth factor (NMF25) were linked with MPPs (Fig3D, E). Next, using this gene module (>150 genes), we performed GSEA with CellMarker data sets and enriched terms indicated that this cluster consists of progenitor cells which could be differentiated towards myeloid or lymphoid lineages (Fig3F), attested that this was indeed MPPs. Other progenitors like HSPCs, CMPs, and CLPs were also distinct (Fig3A, B) and collectively, these results suggest that NMF learns distinct patterns from the data which enables a much better classification of cell types.

**Fig. 3.**
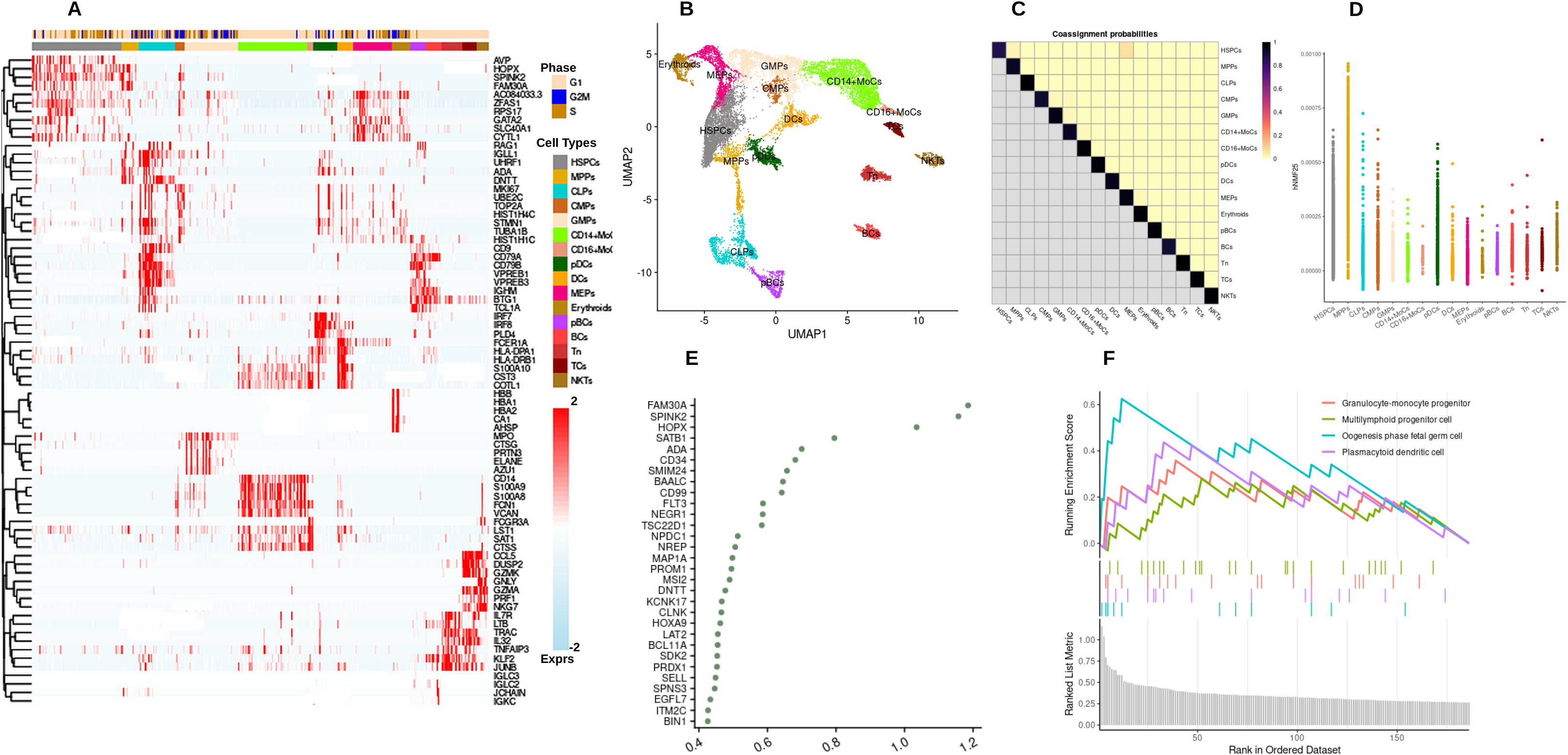
Exploration of Bone marrow SC-data with NMF. **A, Heatmap** representing the top genes, differentially expressed among various cell types. Colored annotation bars represent the cell cycle phase of each cell and cell types classifications. Colored Scales represent a normalized expression. **B, UMAP** representation of each cell class. **C, Heatmap** of coassignmnet probabilities of multiple bootstrap sampling and clustering. **D, Plot** revealing harmony corrected NMF25 separates MPPs from others. **E,F, Top gene** programmes (ranked) linked with MPPs and GSEA based on those modules.

## Discussion

The hematopoietic system is one of the most studied differentiation hierarchies in mammals, and hematopoiesis has long served as a model for how a stem cell maintains and regenerates complex, self-renewing tissues^28^. The advent of single-cell technologies has made us visualize this model in higher resolution but also possesses certain challenges. Single-cell omics datasets are quite high in dimensions and highly sparsed but contain a multitude of transcriptional signals which represent inter and intracellular interactions that control complex biological systems like hematopoiesis^6,10,29^. To infer these interactions, many dimensionality reductions or Matrix factorization (MF; a class of unsupervised techniques) methods are being used frequently (usually PCA). Maximizing the variability captured in specific factors or components is the fundamental concept behind PCA but it may mix the signal from multiple CBPs in a single component. Therefore, PCA may conflate processes and complicate interpretation of the data-driven gene sets from single-cell data or the inference of specific CBPs^10,30^. This is probably the reason when we applied PCA, results indicated that top principal components were able to explain most of the variation in the data, but fails to characterize the progenitor compartments. Although, Tracking the principal components that explain the variations of different progenitors and bootstrapping helped us define a particular class of progenitors (CLPs).

Unlike PCA, ICA extracts hidden factors within data by transforming a set of variables to a new set that is maximally independent (statistically), and theoretically seems a better choice to separate progenitor clusters from single-cell data^10,24,31^. Our results, with ICA, also indicated a better separation among progenitors but few were still hard to differentiate. Again finding the factors associated with different cell classes, and overlapping coassignment probabilities in bootstrap clustering helped us to track other progenitor’s compartments like CMPs. The huge overlappings also suggests the additive patterns of factor recovery from the data with ICA, which indeed could be a better choice when dealing with data sets where cellular hierarchies are the inherent prior biological insights of the data.

NMF decomposes high-dimensional vectors into a lower-dimensional representation. These lower-dimensional vectors and their coefficients are non-negative^10,27^. Theoretically, NMF should also be quite useful for SC datasets as transcriptomic datasets are non-negative in nature^10,32^. Our results with NMF in combination with bootstrapping showed that it recovers distinct gene modules linked with each progenitor compartment, e.g-MPPs which was difficult to classify earlier. Indeed it is probably the best choice to classify progenitors and obviously differentiated lineages as well. In conclusion, here we report the high-resolution maps of human hematopoietic cell fate commitment and lineages via interrogating scRNA transcriptional datasets by using three different matrix factorization (MF) methods (PCA, ICA, and NMF). All the MF method suggests a tri-directional cell developmental trajectory (HSPCs to Erythroids via MEPs, and HSPCs to differentiated myeloid and lymphoid lineages, via different progenitor classes). Our Bootstrap strategy in the combination with gene programs (or CBPs) extracted by the above MF methods was able to differentiate among several progenitor classes as well as recover cellular hierarchies in hematopoiesis. We propose that both ICA and NMF in combination of bootstrapping the clusters are better choices for dimension reduction on SC-data as both find independent sources of variations. ICA can infer additive patterns, good to decide cellular hierarchies, but may have a greater mixture of CBPs. Whereas NMF can find distinctive gene modules from different clusters. As a result, both are better models than PCA, and regardless of the technique selected, the results will also be sensitive to the input data and questions being addressed.

## Methods

### Data Analysis

Single-cell transcriptomic data matrix from BMMCs and CD34^+^ bone marrow cells (obtained from four healthy donors) was retrieved from the GEO repository with accession number GSE139369^33^. Quality control on each SC-data was performed using Matrix R package^34^ and as per the guideline of the broad institute (https://broadinstitute.github.io/2020_scWorkshop/data-wrangling-scrnaseq.html#goal-1).

Seurat V4 pipeline^34^ was used to Normalize, Scale, and variable feature detection from the data in R version 4.1.2. PCA and ICA were performed using *RunPCA* and *RunICA* functions of Seurat, which uses IRLBA^35^ and fastICA^36^ for performing PCA and ICA, respectively. NMF was performed on a log-normalized count matrix using nmf function of an R package RcppML^37^. To correct the donor-specific batch effects, an R package harmony^38^ was used in each case (i.e-PCA, ICA, and NMF). The corrected MFs were later used for running UMAP. The number of clusters were optimized using the clustree R package^39^. Cluster markers were obtained using the recommended negative binomial test with *FindAllMarkers* function in Seurat followed by heatmap visualization using R-packages *ggplot2* and dittoSeq^40,41^. Cell cycle scores were calculated by using the *CellCycleScoring* function from Seurat. We devised a bootstrapping strategy inspired by scran R-package^42^ and usable for Seurat objects. All the codes and instructions to perform bootstrapping are provided on the Github page (https://github.com/PrashINRA/BootStrap_SingleCell). We performed bootstrap sampling and clustering with 100 iterations in each case. GSEA was performed using ranked gene list with ClusterProfiler R package^43^.

## Notes

### Competing Interest Statement

The authors have declared no competing interest.

